# Enhancer activation from transposable elements in extrachromosomal DNA

**DOI:** 10.1101/2024.09.04.611262

**Authors:** Katerina Kraft, Sedona E. Murphy, Matthew G. Jones, Quanming Shi, Aarohi Bhargava-Shah, Christy Luong, King L. Hung, Britney J. He, Rui Li, Seung Kuk Park, Michael T. Montgomery, Natasha E. Weiser, Yanbo Wang, Jens Luebeck, Vineet Bafna, Jef D. Boeke, Paul S. Mischel, Alistair N. Boettiger, Howard Y. Chang

**Author notes:** These authors contributed equally to this work (Katerina Kraft, Sedona E. Murphy, Matthew G. Jones).

## Abstract

Extrachromosomal DNA (ecDNA) drives oncogene amplification and intratumoral heterogeneity in aggressive cancers. While transposable element (TE) reactivation is common in cancer, its role on ecDNA remains unexplored. Here, we map the 3D architecture of *MYC*-amplified ecDNA in colorectal cancer cells and identify 68 ecDNA-interacting elements (EIEs)—genomic loci enriched for TEs that are frequently integrated onto ecDNA. We focus on an L1M4a1#LINE/L1 fragment co-amplified with *MYC*, which functions only in the ecDNA amplified context. Using CRISPR-CATCH, CRISPR interference, and reporter assays, we confirm its presence on ecDNA, enhancer activity, and essentiality for cancer cell fitness. These findings reveal that repetitive elements can be reactivated and co-opted as functional rather than inactive sequences on ecDNA, potentially driving oncogene expression and tumor evolution. Our study uncovers a mechanism by which ecDNA harnesses repetitive elements to shape cancer phenotypes, with implications for diagnosis and therapy.

## Introduction

Extrachromosomal DNA (ecDNA) is a prevalent form of oncogene amplification present in approximately 15% of cancers at diagnosis.^1–5^ EcDNAs are megabase-scale, circular DNA elements lacking centromeric and telomeric sequences and found as distinct foci apart from chromosomal DNA.^6^ Recent work has underscored the importance of ecDNA in tumor initiation and various aspects of tumor progression, such as accelerating intratumoral heterogeneity, genomic dysregulation, and therapeutic resistance.^7–11^ The biogenesis of ecDNA is complex and tied to mechanisms that induce genomic instability, such as chromothripsis and breakage-fusion-bridge cycles, which are prevalent in tumor cells.^6,12–17^

A key aspect of ecDNA function is their ability to hijack *cis*-regulatory elements that increase oncogene expression beyond the constraints imposed by endogenous chromosomal architecture.^18–23^ Consequently, their nuclear organization is tightly tied to their ability to amplify gene expression.^18,20^ Likewise, repetitive genomic elements provide a vast network of cryptic promoters or enhancers capable of re-wiring gene regulatory networks for proto-oncogene expression–including long-range gene regulation.^24–26^ By investigating the 3-dimensional organization of ecDNA, we identified an enrichment of repetitive elements associated with ecDNA structural variation, which we classify as ecDNA-interacting elements (EIEs). We found that insertion of a particular EIE containing a fragment of an ancient L1M4a1 LINE within ecDNA leads to expression of said element that is critical for cancer cell fitness. Our data reveal a relationship between the presence of specific repetitive elements and aberrant expression of oncogenes on ecDNA.

## Results

### ecDNA structural variants enriched for repetitive element insertions

To interrogate the conformational state of ecDNA, we performed Hi-C on COLO320DM colorectal cancer cells (**Fig. 1A)**. Previous investigation of COLO320DM utilizing DNA fluorescent *in situ* hybridization (FISH) and whole-genome sequencing (WGS) identified a highly-rearranged (up to 4.3 MB) ecDNA amplification containing several genes including the oncogene *MYC* and the long non-coding RNA *PVT1*.^18,20^ As a large fraction of the ecDNA in COLO320DM is derived from chromosome 8, with smaller contributions from chromosomes 6, 16, and 13, we elected to focus on the chromosome 8 amplified locus containing *MYC* and *PVT1.*^20^

**Figure 1:**
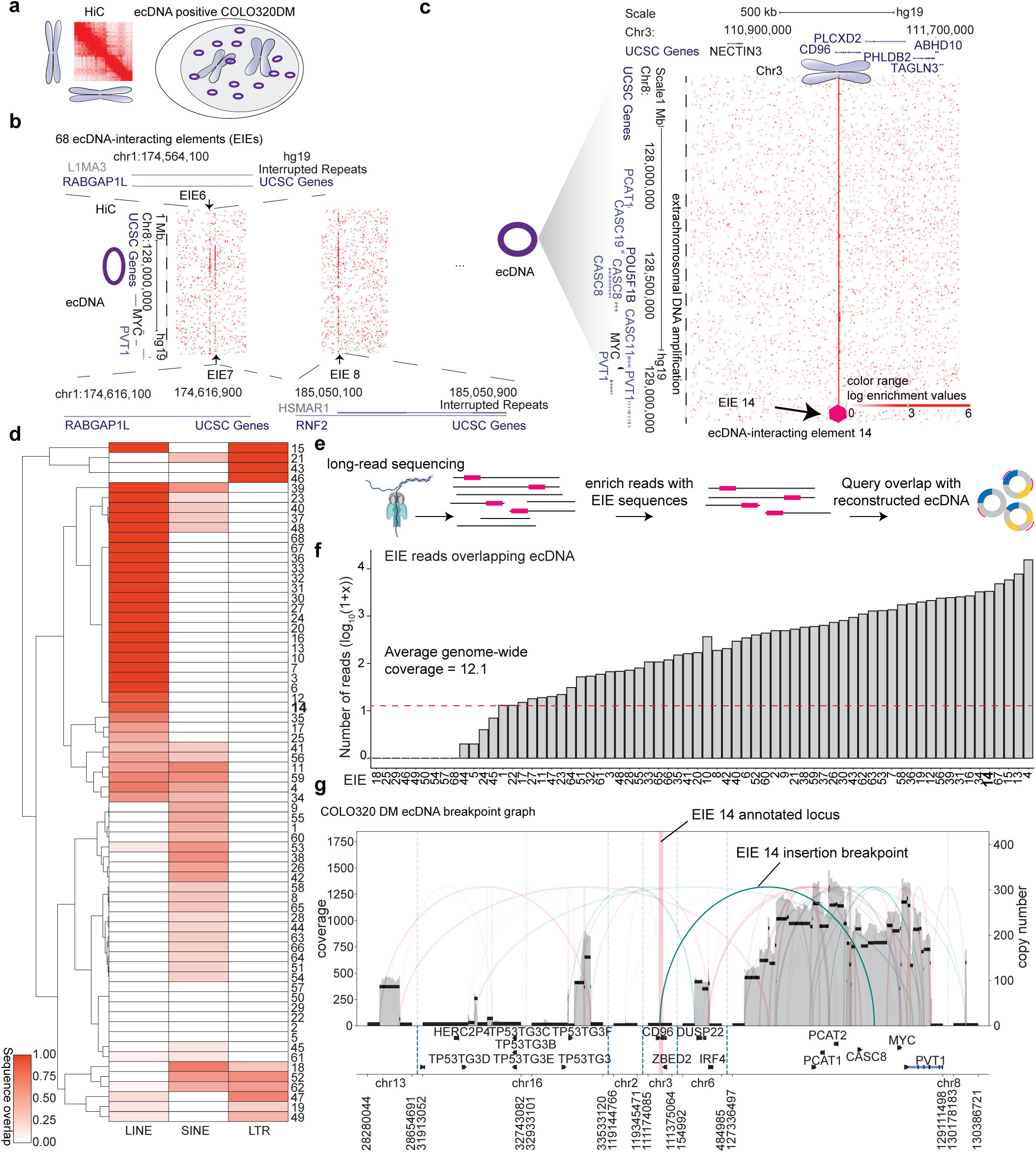
Identification of ecDNA interacting elements (EIEs) **a.** Method schematic of Hi-C performed in the ecDNA containing COLO320DM cell line. **b.** Identification of ecDNA-interacting elements (EIEs). 68 Individual EIEs were manually annotated across all chromosomes based on the interaction across the entirety of the *MYC-* amplified region of chromosome 8. The visualization represents the ecDNA from chromosome 8 with 3 examples of ecDNA-interacting elements (EIEs) localized on other chromosomes. **c.** An example of a specific interaction, EIE 14 on chromosome 3, is enlarged and associated genes are shown for both loci. Arrow and purple hexagon indicate EIE. **d.** Overlap fraction of EIE sequence and annotated LINE, SINE, and LTR elements reported in RepBase. EIEs are clustered according to similarity in overlap fraction across these three classes of repetitive elements. **e.** Pipeline for using Oxford Nanopore ultra-long read sequencing to identify the overlap of ecDNA genomic intervals and EIE-containing reads. **f.** The number of reads that contain a particular EIE and overlap with an ecDNA interval in the COLO320DM cell line. Counts are reported as log_10_(1+x). Average genome coverage (12.1) is represented as a red dashed line. **g.** Reconstruction of the ecDNA breakpoint graph for COLO320DM from Oxford Nanopore ultra-long read data using the CoRAL algorithm. The EIE14 region is highlighted in red and the breakpoint indicating its translocation to the amplified chr8 locus is annotated. Source numerical data are available in source data.

Analysis of the Hi-C maps identified 68 interactions between the chromosome 8 amplified ecDNA locus and other chromosomes that displayed a striking pattern (**Fig. 1B; Supplementary Table T1)**. By binning the data at 1kb resolution, we found that linear elements in the genome contacted the entirety of the megabase-scale ecDNA amplification in a distinctive stripe (**Fig. 1B-C)**. These contacts were spread across all chromosomes in the genome (**Supplementary Table T1**). This atypical interaction pattern suggested a complex structural relationship between the chromosome 8 amplified ecDNA and the endogenous chromosome regions (**Fig. 1B-C**). Further inspection revealed these genomic interactions were enriched for transposable elements annotated as LINEs, SINEs, and LTRs (**Fig. 1D; Extended Data Fig. 1A**, **Supplementary Table T2 and T3**). As these retrotransposons can acquire the ability to regulate transcription when active, we reasoned that the spatial relationship with oncogenes like *MYC* may be important for enhanced expression in COLO320DM cells.^27,28^ We hereafter referred to these 1kb interactions, often containing retrotransposons, as <u>e</u>cDNA interacting <u>e</u>lements (or, EIEs).

While Hi-C is a widely-used method to map genome-wide chromatin interactions, it can be repurposed to identify structural variants, including rearrangements that are a hallmark of cancer genomes.^29,30^ We considered that the atypical striping pattern observed in our Hi-C data was most likely a result of structural variation either in the COLO320 genome or structural variation due to insertion of repetitive elements into ecDNA. To discern between these two possibilities, we performed long-read nanopore sequencing (**Methods**). We chose long-read sequencing to also capture potential heterogeneity in insertion sites in the case of single or multiple integrations (**Fig. 1E-F**; **Methods**). We generated median read lengths of 67,000 bp with the longest read spanning 684,457 bases. Across the 68 EIEs identified, we determined that each participated in a broad spectrum of structural variation - some involved with hundreds or thousands of different rearrangement events (**Extended Data Fig. 1B; Methods)**.

### EIE 14 is a “passenger” on MYC ecDNA

After confirming that the identified EIEs were associated with structural rearrangements, we next investigated the overlap between ecDNA and EIE rearrangements. We first reconstructed ecDNA utilizing the *CoRAL* algorithm^31^, a pipeline that leverages long read data to accurately infer a set of ecDNA from the breakpoints (i.e., structural variation) associated with amplified regions of the genome (**Methods**). We found that reads containing EIEs often overlapped ecDNA intervals at greater coverage than expected from the average genome coverage of our dataset (approximately 12.1), suggesting that these EIEs were contained on at least a subset of ecDNA amplifications **(Figure 1E-F**). We further investigated *CoRAL*’s reconstruction of COLO320DM’s complex and heterogeneous *MYC*-containing amplicon and identified a high-confidence breakpoint connecting a chromosome 3-amplified EIE (EIE14) to an intergenic region between *CASC8* and *MYC* on the chromosome 8 amplification. (**Figure 1G**; **Methods**).

We selected this EIE (EIE 14) for further characterization of EIE biology due to its proximity to *MYC* on the ecDNA and because it contains a segment with homology to L1M4a1, an ancient element distantly related to LINE-1. The percent of nucleotide conservation of this segment to the L1M4a1 consensus sequence is consistent with the L1M4a1’s Kimura divergence value of 34%. We reasoned that this degree of sequence divergence would allow us to specifically target and interrogate its function without unintentionally targeting other repetitive elements in the genome. We also found a fragment of LINE-1 PA2 and an ORF-2 like protein on EIE 14 (**Extended Data Fig. 1C-D**).^32,33^ Although the mechanism generating the adjacency of the fragments remains uncertain, the L1M4a1-like segment harbors a polyA-signal–like motif (AAAAAG), supporting a model in which an L1PA2 transcript read through its own 3′ end and terminated at this neighboring signal, producing a 3′-transduced RNA that could be mobilized in trans by LINE-1 enzyme (**Extended Data Fig. 1C-D**).^32,33^

To confirm the computational reconstruction of the ecDNA and the heterogeneity of different ecDNA molecules, we turned to CRISPR-CATCH - a method for isolating and sequencing ecDNA - to elucidate the size and variations of ecDNAs containing EIE 14 (**Fig. 2A**).^22^ Targeting EIE 14 with two independent gRNAs, we successfully isolated ecDNA fragments from the COLO320DM cell line for sequencing (**Fig. 2B**). Sequence analysis of these bands confirmed the presence of EIE 14, originally annotated on chromosome 3, to be inserted onto chromosome 8 between the *CASC8* and *CASC11* genes approximately 200 kilobases away from *MYC*, in agreement with the long-read nanopore sequencing (**Fig. 1G**, **Fig. 2C, Extended Data** Fig.2A **& B and Supplementary Table T4-T6).** Multiple bands of different sizes on the PFGE gel indicated the presence of varying sizes of ecDNAs, all sharing the EIE 14 insertion within the chromosome 8 amplicon **(Fig. 2B-C)**. Beyond EIE 14, the CRISPR-CATCH approach allowed us to capture and sequence a subset of EIEs initially identified through Hi-C analysis (**Fig. 2D**). The identification of the additional EIEs observed in the Hi-C data suggest that the “striping” between the ecDNA and endogenous chromosomes is an artifact of these sequences’ presence on ecDNAs, rather than true *trans* contacts, at least for this identified subset. Though the recent T2T genome build^34^ annotates EIE 14 to chromosome 3 (**Extended Data Fig. 2C)**, we found evidence that the structural variant described here between EIE 14 and the *MYC*-containing amplicon region is identified as a translocation event between Chr8:128,533,830 and Chr3:111,274,086 in approximately 46% (minor allele frequency of 0.467646) of non-disease individuals (**Supplementary Table T4 (row 7)**).^35^ This suggests that this structural variant was pre-existing prior to cancer formation in the COLO320-originating patient and was subsequently amplified as a passenger on ecDNA.

**Figure 2:**
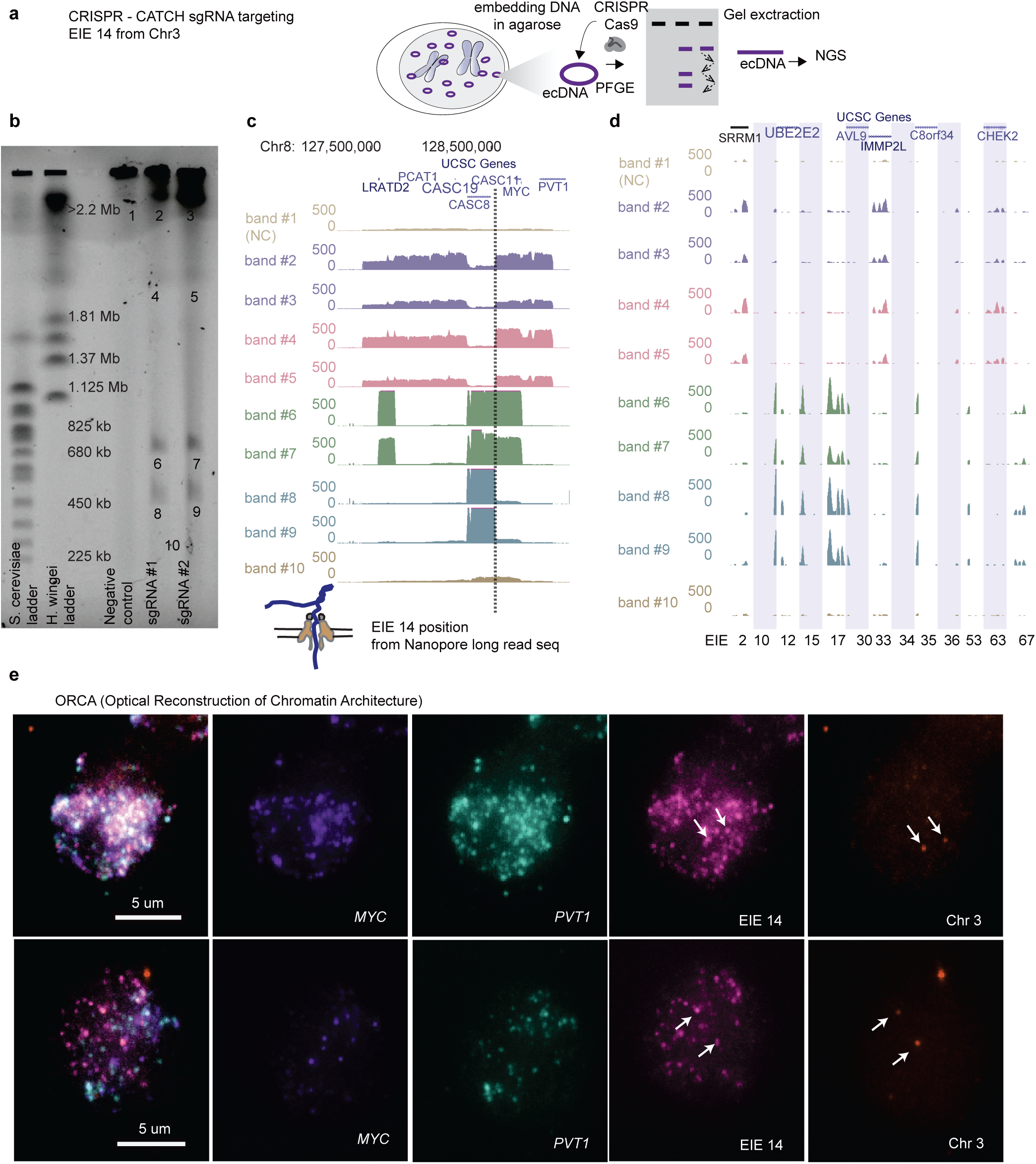
CRISPR-CATCH Elucidates ecDNA Composition and EIE Insertions. **a.** Schematic diagram illustrating the CRISPR-CATCH experiment designed to isolate and characterize ecDNA components. The process involves the use of guide RNA targeting the EIE 14 from chromosome 3. DNA is embedded in agarose, followed by pulse-field gel electrophoresis (PFGE), allowing for the band extraction and subsequent next-generation sequencing (NGS) of ecDNA fragments. **b.** The PFGE gel image displays the separation of DNA fragments, lines from left ladder, ladder, empty lane, Negative control, sgRNA #1, sgRNA #2 and band numbers for NGS seen in **C-D**. EIE 14 targeted by the guide RNAs leads to cutting of the ecDNA’s chromosome 8 sequences to form multiple discrete bands, confirming EIE 14 insertion onto ecDNA. sgRNA #1 ATATAGGACAGTATCAAGTA; sgRNA #2 ATATTATTAGTCTGCTGAA; Full EIE 14 sequences from long-read sequencing is in Supplementary Table T6. **c.** Whole genome sequencing results confirm the presence of EIE 14, originally annotated on chromosome 3, within the ecDNA, between the *CASC8* and *CASC11* genes, approximately 200 kilobases upstream from *MYC*. The dotted line indicates the position of this insertion. Each band is an ecDNA molecule of a different size that contains the EIE 14 insertion. **d.** Additional EIEs identified in the initial Hi-C screen, captured, and sequenced in the CRISPR-CATCH gel bands from (**B**), each EIE is one one vertical shaded box with coordinates and denote insertion events within the ecDNA. **e.** ORCA (Optical Reconstruction of Chromatin Architecture) visualization of the COLO320DM cell nucleus. The max-projected images show the spatial arrangement of the *MYC* oncogene, EIE 14 and the *PVT1* locus, labeled in different colors for two different cells. Left most panel is an overlay of all images registered to *nm* precision (see **Methods**).The scale bar represents 5 micrometers. Chr3 probe maps to the breakpoints of the EIE 14 origin inside *CD96* intron. Source numerical data and unprocessed blot are available in source data.

### EIE 14 makes frequent contact with MYC

We then utilized Optical Reconstruction of Chromatin Architecture (ORCA) to quantify the spatial relationship of EIE 14 with *MYC* (**Fig. 2E**). ^36,37^ Barcoded probes were designed targeting the unique portion of EIE 14 (1kb), *MYC* exon 2 (3.1kb), *PVT1* exon 1 (2.5kb), and the endogenous chromosome 3 region flanking of EIE 14 (3kb) **(Supplementary Table T7**) to determine the spatial organization of EIE 14 relative to the ecDNA. These specific exons were chosen to account for the fact that amplicon reconstruction of ecDNA in the COLO320DM cell line demonstrated an occasional rearrangement of *MYC* exon 2 replacement by *PVT1* exon 1.^20^ Since EIE 14 is classified as a repetitive element, we confirmed probe specificity by staining the EIE 14 locus in K562 cells that do not contain ecDNA. Indeed, we detect only 1-3 labeled regions in the non-amplified context (**Extended Data Fig. 3A**). In contrast, when labeling COLO320DM cells, EIE 14 colocalized with the ecDNA and amplified to a similar copy number per cell (**Fig. 2E, Extended Data Fig. 3B**). The extensive structural variation detected in the long-read sequencing and the amplification of EIE 14 visualized by ORCA (**Extended Data Fig. 3B**) suggest a model where the element resides in the sequence amplified on ecDNA and participates in *cis* and/or *trans-*contacts with other ecDNA molecules.

It has been proposed that amplified loci within ecDNA are able to regulate oncogene expression through *cis*-interactions on the same ecDNA molecule as well as *trans*-interactions between ecDNAs via a clustering mechanism.^20^ As such it is important to understand not only the structural variations of ecDNA, but also how they are arranged in the nucleus for a comprehensive understanding of potential regulatory function. We quantified the spatial distributions of *MYC* exon 2, *PVT1* exon 1, and EIE 14; the imaged loci were fitted in 3-dimensions with a gaussian fitting algorithm to extract x,y,z coordinates (**Fig. 3A-C, Methods**). The copy number of identified loci varied from zero detected points to 150 per cell. On average, *MYC* had 29, *PVT1* had 31 and EIE 14 had 22 copies per cell (**Extended Data Fig. 3B**). Similar distributions of points-per-cell, as well as strong correlation (*r*>0.7) between number of points per loci per cell (**Extended Data Fig. 3C)** suggests that this EIE is not inserted into multiple sites on a single ecDNA.

**Figure 3:**
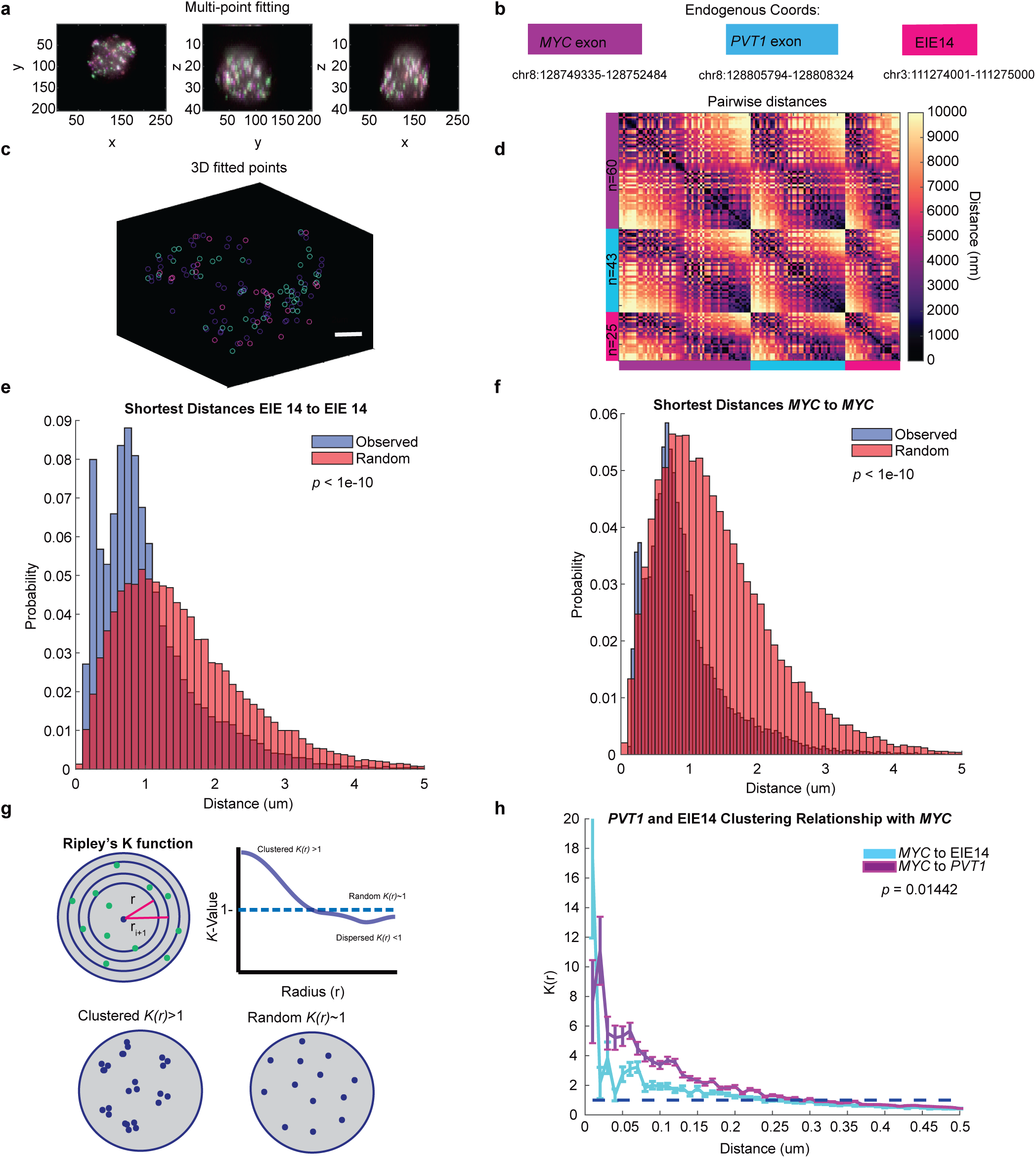
EIE 14 spatially clusters with *MYC*. **a.** X, Y, Z projections of *MYC* exon (purple), *PVT1* (blue), and EIE 14 (pink) **b.** Endogenous coordinates of all three measured genomic regions. **c.** Single cell projection of the 3D fitted points from (**A**). **d.** Pairwise distances between *MYC* (purple), *PVT1* (blue), and EIE 14 (pink) of a single cell. Number of fitted points per genomic region *n=60, n=43,* and *n=25* respectively. **e.** Histogram of distribution of distances of the observed shortest pairwise EIE 14 to EIE 14 distances and the expected shortest pairwise distances of points randomly simulated in a sphere (two-tailed Wilcoxon *ranksum p<*1e-10) of *n=*1329 analyzed cells across 2 biological replicates. **f.** As in (**E**) but for *MYC* to *MYC* shortest pairwise distances (Two-tailed Wilcoxon *ranksum p<*1e-10). **g.** Schematic of Ripley’s *K* function to describe clustering behaviors over different nucleus volumes. Top shows the nucleus divided into different shell intervals and how the *K* value is plotted for increasing radius (*r).* Bottom shows an example of what clustered *K(r)>1* vs. random *K(r)∼1* points could look like. *K-*values greater than one indicate clustering behavior relative to a random distribution over that given distance interval (*r), K* values ∼ one denote random distribution, while *K* values less than one indicate dispersion behavior **h.** The average *K(r)* value across distance intervals of 0.01 to 0.5 um in 0.02 um step sizes to describe the clustering relationship of *PVT1* and EIE 14 relative to *MYC* across different distance intervals (um). Error bars denote SEM. (Two-tailed Wilcoxon *ranksum p=*0.01442). Source numerical data are available in source data.

Once the centroids of each point per cell were identified (**Fig. 3C**) we calculated the all-to-all pairwise distance relationship (**Fig. 3D**). The off-diagonal pattern of distances between EIE 14, *MYC*, and *PVT1* suggested a tendency for these loci to cluster at genomic distances <1000nm. We further quantified the spatial relationships across all 1329 imaged cells by calculating the shortest pairwise distances between the three loci. To determine if these ecDNA molecules were spatially clustering in cells, we leveraged our observation that each ecDNA molecule carries a single copy of *MYC* and *EIE 14*. Thus, distances between *MYC* and other *MYC* loci should be closer than random if the ecDNA were spatially clustered. Random distances were simulated in a sphere with the identical number of points per a given cell. The distribution of shortest pairwise distances between *MYC* and *MYC* and between *EIE 14* and *EIE 14* were left-shifted compared to the randomly simulated points, suggesting a nonrandom organization (**Fig. 3E-F**, *p*<1e-10). The median observed versus expected distances between each *EIE 14* loci were 748 nm and 927 nm respectively and the median observed versus expected distances between each *MYC* loci were 707nm and 814nm respectively.

Previous work has proposed that enhancers can exert transcriptional regulation on promoters at a distance of up to 300 nm via accumulation of activating factors.^38–41^ To determine whether EIE 14 and *MYC* are within this regulatory distance range on ecDNA molecules, we calculated the pairwise distances between loci. Though the median distances between *MYC* and EIE 14 (797 nm) and *PVT1* (585 nm) were greater than 300 nm, 12% and 20% of these loci, respectively, were within the regulatory range of *MYC* (**Extended Data Fig. 3D-E**).

To investigate the spatial relationship between EIE 14 and *MYC* while controlling for locus density, we calculated the degree of spatial clustering across distance intervals using Ripley’s *K* spatial point pattern analysis (See **Method**s, **Fig. 3G**)*. MYC* exhibited the strongest clustering with *EIE 14* at distances less than 200 nm (K-value > 1), and this behavior approached a random distribution at greater distances (K-value ∼ 1; **Fig. 3G,H**). While, on average, distances between *MYC* and EIE 14 were further than *MYC* and *PVT1 (***Extended Data Fig. 3D-E,** at distances <300 nm EIE 14 and *PVT1* displays a similar clustering behavior with *MYC* **Fig. 3H**). This clustering suggests that EIE 14 is acting as a proximity-dependent regulator of *MYC* reminiscent of enhancer-promoter interactions.^42^ Altogether, the spatial clustering behavior of this ecDNA species measured here and previously^20^, the propensity for *MYC* to engage in “enhancer hijacking”^43^, and the ability of reactivated repetitive elements to engage in long-range gene activation^27^ suggests that any genomically linear separation of *MYC* and EIE 14 is overcome in both *cis-* (interaction with *MYC* on the same ecDNA molecule) and *trans* (ecDNA-ecDNA interactions).

### EIE 14 is critical for cancer cell fitness and displays enhancer activity

To test whether the identified transposable elements are important for the cancer cell proliferation, we performed a CRISPR interference (CRISPRi) growth screen targeting a subset of EIEs in COLO320DM cells engineered to stably express dCas9-KRAB(**Fig. 4A-B)**.^46^ We were able to target 36 out of the 68 EIEs with sgRNAs that met the following criteria: (1) must meet stringent specificity criteria to reduce potential off targets intrinsic to repetitive sequences (see **Methods**) and (2) have at least two sgRNAs per EIE. We also included 125 non-targeting controls (NTC) that were introduced into cells with the EIE sgRNAs via lentiviral transduction (**Supplementary Table T10**). Post-transduction, we monitored cell proliferation at multiple time points: 4 days (baseline), 3 days after baseline, 14 days, and 1 month (30 days), followed by deep sequencing to quantify sgRNA frequencies (**Fig. 4B)**. We obtained highly reproducible guide counts across replicates and timepoints(**Extended Data Fig. 4B-C**).

**Figure 4:**
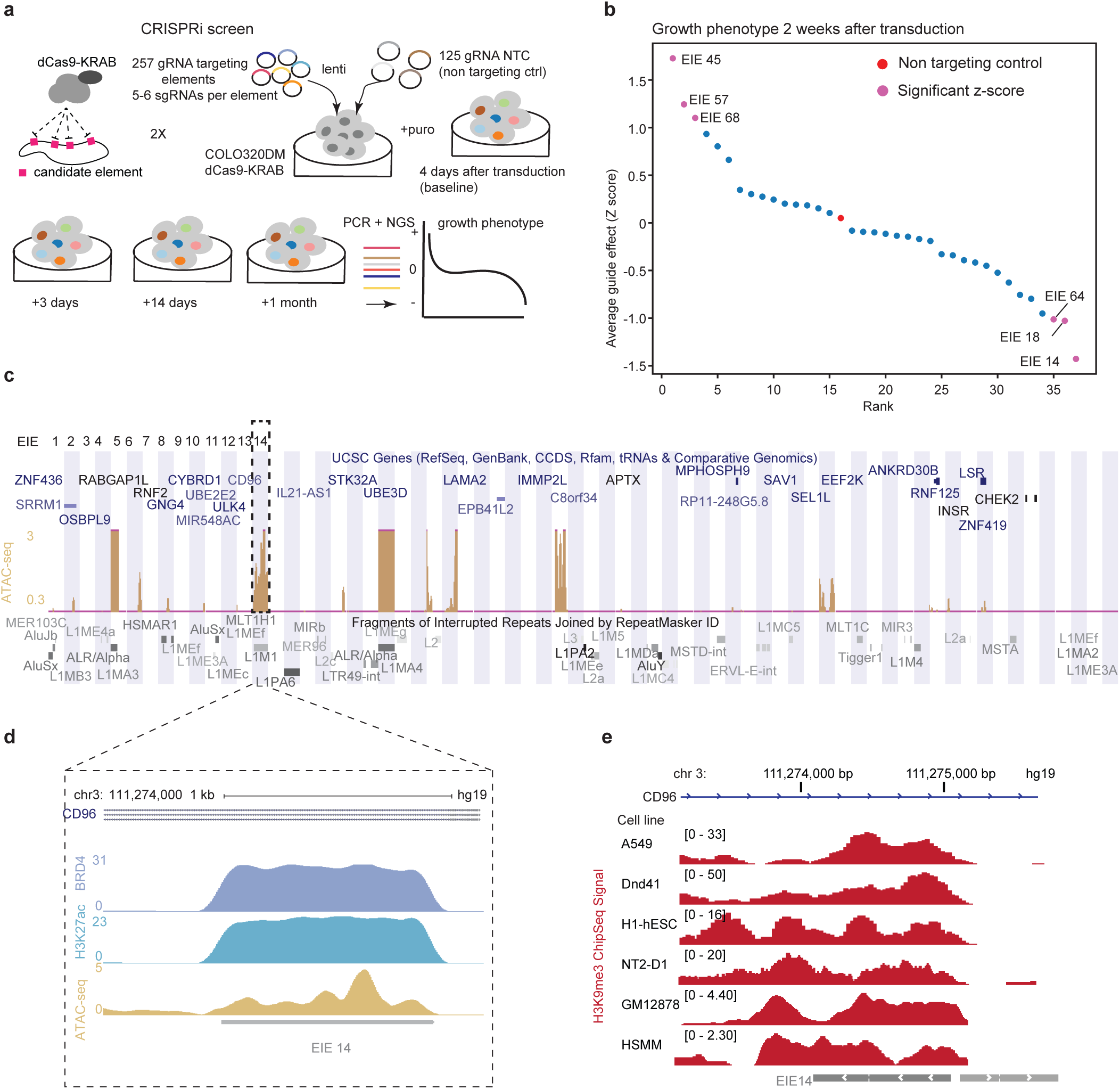
EIE 14 is important for cell proliferation and has enhancer signatures. **a.** Schematic of the CRISPRi screening strategy used to evaluate the regulatory potential of the 68 EIEs by designing 4-6 gRNAs per element for a total of 257 genomic regions tested and 125 non-targeting control sgRNAs. The screen involved the transduction of cells with a lentivirus expressing dCas9-KRAB and the sgRNAs such that each cell received 1 sgRNA, followed by calculation of cell growth phenotype over a series of time points (Baseline(4 days), Baseline + 3 days, Baseline + 14 days, and Baseline + 1 month). The screen was further filtered on guide specificity (**methods**) and 36/68 targeted EIEs met the qualifying threshold. **b.** The growth phenotype of COLO320DM cells 2 weeks post-transduction, relative to non-targeting control (NTC). Each point represents the average guide effect (Z-score) for sgRNAs targeting the 36 qualifying EIEs, ranked by their impact on cell growth. EIE 14 is indicated by dashed rectangle with negative Z-score < -1 (significant negative impact on cell viability). See Extended Data for additional timepoints. Positive hits are labeled in pink with their corresponding EIE. **c.** UCSC Genome Browser multi-region view showing the locations of the EIEs within the genome. Each EIE is indicated by a vertical bar. The browser displays the annotations for genes and repetitive elements such as *Alu*, LINE, and LTR elements (RepeatMasker), ATAC-seq dataset^20^ is normalized for copy number (see **Methods**). **d.** Zoom-in of EIE 14’s histone marks: enrichment of H3K27 acetylation^18^, BRD4 binding^20^, and ATAC-seq peaks. ChIP data was normalized to input to control for copy number. ATAC-seq data was normalized to library size (**methods**). **e.** H3K9me3 histone modification of EIE 14 across ENCODE cell lines.^44,45^

Our data showed that the growth phenotype curve for three out of thirty six of our targeted EIEs at various time points indicated a Z-score of less than -1, which suggested a significant negative impact on cell viability, with an acute growth defect after only 3 days (**Fig. 4B, Extended Data Fig. 4, Supplementary Tables T8 and T9)**. These elements were categorized as evolutionarily older based on their retrotransposition activity in the human genome and spanned classes (LINEs, SINEs, LTRs) (**Supplementary Table 11**). The enrichment of old TEs may be confounded by the relative ease of targeting sequences with increased sequence divergence. They are generally found in gene poor regions making it unlikely that silencing would lead to secondary effects from heterochromatin spreading. Collectively, these results suggest that a subset of our targeted EIEs, including EIE 14, can contribute to cancer cell growth and fitness. We speculate that this is related to EIE interaction with *MYC*, as knockdown of this oncogene has been shown to have similar effects on COLO320DM growth and survival.^47,48^ Additionally, three out of thirty six of the measured EIEs also had a Z-score greater than 1, indicating a significant increase of cell growth or fitness. The identity of these elements also spanned element classes with two (EIE 68 and EIE 45) being located within two uncharacterized ncRNAs and one (EIE 57) within the first exon of the ANKRD30B protein coding gene which has been implicated in cell proliferation.^49^ Further investigation of these hits are warranted in future studies to explain their positive effects on cell growth, especially those within the uncharacterized ncRNA regions.

The strongest growth defect was observed for perturbation of EIE 14 (**Fig. 4B)**, which when combined with our finding of its co-localization with ecDNA-amplified *MYC* (**Fig. 3H**), suggests a potential enhancer-like regulatory role for this EIE. To examine the epigenetic landscape of this element we leveraged copy-number normalized ChIP-seq measuring H3 lysine 27 acetylation (H3K27ac), BRD4 occupancy, and ATAC-seq accessibility data. These epigenetic features are all commonly associated with enhancer activity.^18,50,51^ Notably, many EIEs, including EIE14 were accessible in COLO320DM cells (**Fig. 4C-D, Extended Data Fig. 5**). The measured accessibility of EIE 14 contrasts the normally silenced H3 lysine 9 trimethylation (H3k9me3) state across annotated human cell lines (**Fig. 4E**).^44,45^ Cross-referencing our identified EIEs with accessibility data from other ecDNA containing cell lines demonstrated that accessibility of EIEs is a more generalizable phenomenon beyond COLO320DM cells (**Extended Data Fig. 5**). Altogether, the accessibility and proximal clustering of EIE14 points towards active regulatory potential of this element in COLO320DM cells, while identification of accessible EIEs across cell lines suggests a broader functional relevance of EIE regulatory potential on ecDNA (**Extended Data Fig. 5)**.^50,51^

To determine whether EIE 14 activity is a consequence of ecDNA formation, we performed RNA-FISH on the sequence-specific 1kb segment of EIE 14 in COLO320DM and isogenic COLO320HSR cells. The HSR or homogeneously staining region cell line contains a similar copy number amplification of the *MYC*-amplified portion of chromosome 8, but the majority of these copies have integrated into chromosomes (**Fig. 5A**).^18^ We reasoned that if the unique extrachromosomal context of ecDNA facilitates activation of EIE 14, we should not see evidence of its activity in the COLO320HSR genome-integrated context. Indeed, we observed distinct transcription events in the DM line (median *n=*8 transcripts per cell) but not in the HSR line(median *n=*0 transcripts per cell; **Fig. 5B, Extended Data Fig. 6A-B).**

**Figure 5:**
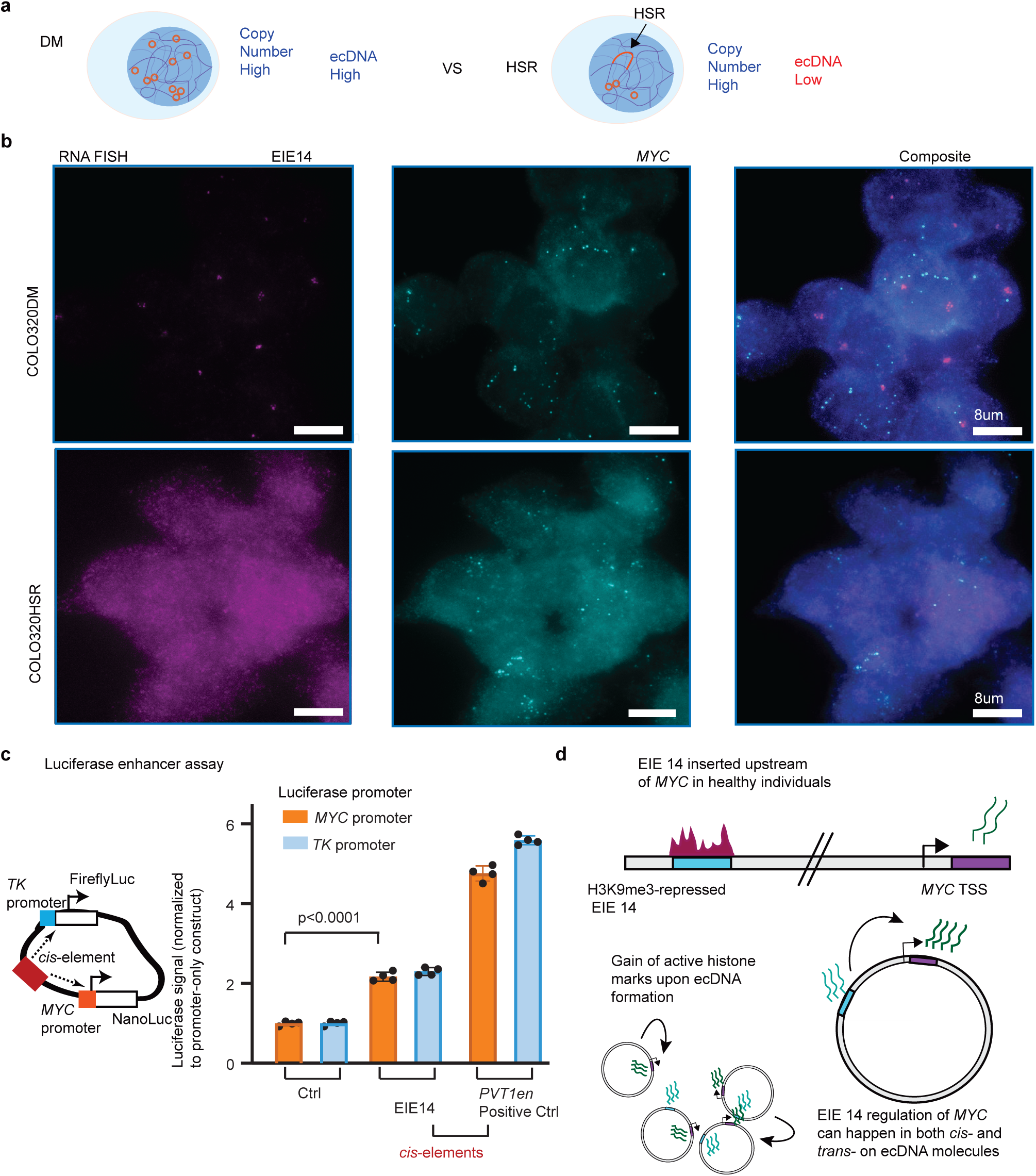
ecDNA context is critical for EIE 14 enhancer activity. **a.** (Top) Schematic outlining COLO320DM cell line as high copy number and high ecDNA vs HSR-as high copy number but low ecDNA. **b.** RNA-FISH labeling for EIE 14 and *MYC* exon 2 transcription in COLO320 DM and HSR. Median transcripts for EIE 14 are 4 and 0 for the DM and HSR cells ( two-tailed wilcoxon ranksum *p=*8.22 10-94), respectively. DM cells have a median of 14 *MYC* transcripts and HSR cells have a median of 8 transcripts per cell (two-tailed wilcoxon ranksum *p=*2.18 10-66). *n=*712 cells (DM) *n=*681 (HSR) across 2 biological replicates. **c.** Luciferase enhancer assay schematics and fold change in luciferase signal driven by either *MYC* or TK promoter normalized to promoter-only construct. *n=*4 biological replicates. EIE 14 compared to positive control (*PVT1* positive control from^20^). *P-*values obtained from two-tailed unpaired t-test. Error bars are standard deviations from the mean. **d.** Schematic outlining EIE 14 as a translocation event in healthy patients where EIE 14 is normally inactive across annotated cell lines (Fig. 5A). EIE 14 gains regulatory potential when it is amplified within ecDNA as a consequence of translocation near *MYC.* EIE 14 can then act as a regulator of *MYC* in both *cis-* and *trans-*contacts within and between ecDNAs. Source numerical data and images are available in source data.

Finally, to directly test the ability for the EIE 14 sequence to act as an enhancer of *MYC* expression, we performed a luciferase reporter assay measuring its ability to activate transcription *TK* and *MYC* promoters **(Fig. 5C).**^20,52^ EIE 14 significantly increased *MYC* promoter-mediated reporter gene expression relative to the promoter only control, signifying bona fide enhancer activity (**Fig. 5C**). Separating EIE 14 into L1M4a1 and L1PA2 fragments further demonstrated that both sequences can individually act as enhancers, with an additive effect when combined (**Extended Data Fig. 6C**). In sum, the enhancer-associated features and regulatory activity of the luciferase assay suggested that EIE 14, and possibly other EIEs, have been co-opted as regulatory sequences when found on ecDNA, influencing the expression of ecDNA-borne oncogenes (**Fig. 5D**).

## Discussion

This study uncovers a mechanism by which transposable elements (TEs), typically silenced by heterochromatin, may acquire regulatory potential when amplified on extrachromosomal DNA (ecDNA).^53–55^ Somatically active retrotransposition events^56^ as induced by LINEs and SINEs, are abundant in the human genome and represent a major source of genetic variation.^57^ Across cancer types, retrotransposon insertions contribute significantly to structural variation, genomic rearrangements, copy number alterations, and mutations—including in colorectal cancer.^58–65^ The activity of these elements in cancer can induce genomic instability and drive the acquisition of malignant traits. For instance, when reactivated LINE-1 elements are inserted into the APC tumor suppressor gene in colorectal cancer, they disrupt gene function and confer a selective advantage.^66^ In other contexts, TEs act as bona fide transcriptional enhancers, amplifying oncogenic gene expression and promoting tumorigenesis.^67^

Here, we describe the enhancer-like activity of a specific identified element, EIE 14, which becomes active through its association with ecDNA (**Fig. 5D**). EcDNAs, which are randomly segregated during cell division, are subject to strong selective pressure.^10^ The recurrent co-amplification of TEs on ecDNA-containing cell lines suggests they may contribute to ecDNA fitness and oncogenic function. We show that retrotransposons like L1M4a1/EIE 14 can escape the inactive chromatin environment of their native genomic loci when inserted within the transcriptionally permissive landscape of ecDNA.^18^ In fact, we demonstrate that EIE 14 is only transcriptionally active in the context of ecDNA and not in the endogenous chromosomal context of the copy-number matched, isogenic COLO320 HSR cells. The context-specific transcription suggests a purely epigenetic regulation imbued by the local environment of ecDNA. This environment enables EIE 14 to potentially influence nearby oncogenes such as *MYC*. Given that LINEs have been shown to exhibit enhancer-like behavior when reactivated,^27,28,68^ the clustering of ecDNA molecules observed through ORCA may further enhance spatial feedback^69^ of both *cis*- and *trans*-regulatory interactions of EIE 14 with oncogenic targets.

Although EIE 14 is incapable of autonomous transposition and lacks a complete L1M4a1 sequence, its subsequent activity upon integration into ecDNA suggest that degenerate ancient sequences may become functionally active under the right conditions. Previous work has shown that single nucleotide polymorphisms associated with familial cancer risk often affect the biochemical activity of noncoding enhancer elements linked to oncogenes activated in cancer.^70,71^ Our results extend this model by proposing that inherited variation in ancient TE insertions, such as EIE 14 near *MYC*, can create latent enhancers that become activated when the oncogene locus is excised into ecDNA.

Perturbation of EIE 14 through CRISPRi resulted in impaired cell growth in COLO320DM cells, indicating that its reactivation contributes to the colorectal cancer phenotype. Quantifying the precise downregulation of *MYC* is constrained by ecDNA heterogeneity, a narrow temporal window in MYC-addicted cells, rapid growth arrest and subsequent loss of successfully targeted cells. While this functional evidence supports a potential oncogenic role, further studies focusing on *in vivo* analyses are necessary to determine whether TEs on ecDNA are sufficient to confer a survival advantage or correlate with poor patient prognosis. Notably, recurrent LINE-1 amplification on ecDNA have been observed in primary esophageal cancer, providing *in vivo* support for the clinical relevance of this phenomenon.^72^

Finally, the amplification of retrotransposable elements onto ecDNA introduces a mechanism for increasing ecDNA structural variation, leveraging the 40% of the genome composed of typically silenced repetitive elements. Retrotranspositions are, in fact, the second-most frequent type of structural variant in colorectal adenocarcinomas.^73^ Just as transposons have played a major role in bacterial plasmid evolution through cycles of insertion and recombination,^74^ our findings allude to a parallel evolutionary trajectory in human oncogenic ecDNAs. The transcriptionally permissive state of ecDNA enables these elements to potentiate oncogene activation and selection—making them both prognostic biomarkers and potential therapeutic targets.

## Acknowledgements

This project was supported by Cancer Grand Challenges CGCSDF-2021\100007 with support from Cancer Research UK and the National Cancer Institute (H.Y.C., P.S.M.), NIH award U01DK127419 (A.N.B.) and NSF grant EF2022182 (A.N.B). M.G.J. is supported by NIH K99CA286968. S.E.M. was supported by a Stanford Bio-X SIGF Fellowship and the NIH award DP5OD037361 through the OD and the NIDCR. A.B.-S. was supported by the Stanford Medical Scholars Research Program and an Alpha Omega Alpha Carolyn L. Kuckein Student Research Fellowship. K.L.H. was supported by a Stanford Graduate Fellowship and an NCI Predoctoral to Postdoctoral Fellow Transition Award (NIH F99CA274692 and K00CA274692). M.T.M. was supported by an NSF Graduate Research Fellowship (DGE-1656518). Y.W. was supported by the Schmidt Science Fellows program. H.Y.C. was an Investigator of the Howard Hughes Medical Institute.V.B. was supported in part by the Cancer Grand Challenges partnership funded by Cancer Research UK (CGCATF-2021/100025) and the National Cancer Institute (OT2CA278635), U24CA264379, and by R01GM114362. We thank Mervinaz Koska for help with luciferase measurement.

## Author Contributions

K.K. and H.Y.C. conceived the project. K.K., S.E.M., M.G.J., and H.Y.C. wrote the manuscript with input from all authors. K.K., S.E.M., Q.S., A.B.S., B.J.H., R.L., N.E.W., and Y.W. performed experiments, M.G.J., S.E.M., C.L., K.L.H., S.K.P., J.L., M.T.M., analyzed the data. J.D.B provided guidance on transposable element analysis. V.B. and P.S.M provided guidance on manuscript content. A.N.B. contributed instrument time and software and advised on image data analysis.

## Methods

### Cell culture

Cell lines were obtained from ATCC. COLO320DM (CCL-220) and COLO320-HSR (CCL-220.1) cells were maintained in RPMI; Life Technologies, Cat# 11875-119 supplemented with 10% fetal bovine serum (FBS; Hyclone, Cat# SH30396.03) and 1% penicillin-streptomycin (pen-strep; Thermo Fisher, Cat# 15140-122). All cell lines were routinely tested for mycoplasma contamination. Presence of ecDNA in cell lines was confirmed via metaphase spreads.

### Hi-C

Ten million cells were fixed in 1% formaldehyde in aliquots of one million cells each for 10 minutes at room temperature and combined after fixation. We performed the Hi-C assay following a standard protocol to investigate chromatin interactions within colorectal cancer cells.^1^ HiC libraries were sequenced on an Illumina HiSeq 4000 with paired-end 75 bp read lengths. Paired-end HiC reads were aligned to hg19 genome with the HiC-Pro pipeline.^2^ Pipeline was set to default and set to assign reads to DpnII restriction fragments and filter for valid pairs. The data was then binned to generate raw contact maps which then underwent ICE normalization to remove biases. HiCCUPS function in Juicer^3^ was then used to call high confidence loops. Visualization was done using Juicebox https://aidenlab.org/juicebox/

### Analysis of EIEs for repetitive element overlap

To assess the overlap of classes of repetitive elements with our identified EIEs, we obtained the “RepeatMasker” and “Interrupted Repeats” tracks from UCSC Genome Browser for hg19. For each EIE, we computed the fraction of the sequence that overlapped with the merged BED file containing the RepeatMasker and Interreputed Repeats annotations. We report the overlap separately for LINE, SINE, and LTR repetitive element classes. Importantly, each EIE is exactly 1kb long so no length normalization is performed. To compute an expected proportion, we computed the fraction of hg19 covered by each repetitive element class. The results are reported in **Figure 1D** and **Extended Data** Figure 1A.

### Whole Genome Sequencing (WGS) with Oxford Nanopore

High-molecular weight (HMW) genomic DNA was extracted from approximately 6 million COLO320DM cells using the Monarch HMW DNA Extraction Kit for Tissue (NEB #T3060L) following the Oxford Nanopore Ultra-Long DNA Sequencing Kit V14 protocol. After extracting HMW gDNA, we constructed Nanopore libraries using the Oxford Nanopore Ultra-Long DNA Sequencing Kit V14 (SQK-ULK114) kit according to manufacturer’s instructions. We sequenced libraries on an Oxford Nanopore PromethION using a 10.4.1. Flow Cell (FLO-PRO114M) according to manufacturer’s instructions. Basecalls from raw POD5 files were computed using Dorado (v.0.2.4).

### Identifying, re-mapping EIE-containing reads, and detecting structural variants

We first identified Nanopore reads containing a single element by aligning reads with minimap2^4^ and filtered out reads that were not mapped by the algorithm (denoted by “*” in the RNAME column of the BAM entry). Then, taking these reads we performed genomic alignment once again using minimap2 against hg19.

From these new alignments of only the reads found to contain the element under consideration, we performed two analyses for each element. First, we detected structural variant detection using Sniffles2.^5^ Second, we identified overlap of reads with ecDNA-containing intervals that were reconstructed with long reads (see section “**Reconstruction of ecDNA amplicons with long-read data”**). In this second analysis (presented in **Figure 1F**), we counted the number of reads covering regions contained with cycles reconstructed with CoRAL algorithm.^6^While this analysis does not explicitly account for reads that originate from chromosomal or extrachromosomal regions, we reasoned that elements that were carried on ecDNA would be amplified and thus these elements would be highly covered; on the other hand, regions that were were primarily chromosomal would be represented by a similar number of reads to the overall genome coverage.

### Reconstruction of ecDNA amplicons with long-read data

We reconstructed ecDNA amplicons from ultra-long Oxford Nanopore reads using the CoRAL algorithm.^6^ Briefly, this algorithm determines focally amplified regions of the genome using CNVkit^7^ and then finds reads that support this focally amplified region. In doing so, CoRAL identifies genomic breakpoints between the focally amplified seed region and disparate parts of the genome to create a “breakpoint graph”. From this breakpoint graph, putative ecDNA cycles are identified. We report the breakpoint graph in **Figure 1G** which includes a breakpoint between EIE14 (annotated on chr3) and an intergenic region between *CASC8* and *MYC* on chr8.

In addition to detecting EIE14 on the *MYC*-amplifying ecDNA in COLO320DM, we additionally quantified the number of reads that span a given EIE and any part of the COLO320DM genome amplified as ecDNA. We report the number reads that support an EIE as amplified on ecDNA in **Figure 1F**.

In **Extended Data** Figure 2B we visualized reads connecting EIE14 on chr3 with the chr8 ecDNA-amplified region using Ribbon (v 2.0.0).^8^

### ATAC-seq analysis and normalization

ATAC-seq and ChIP-seq data for COLO320DM and SNU16 was obtained from Hung, Yost *et. al.* 2021^9^ and for PC3 and GBM39KT from Wu *et. al.* 2019^10^. Previously, ATAC-seq data was mapped to hg19. While ChIP-seq data was normalized to input, as input is not sequenced with ATAC-seq, these data were further normalized by library size. Specifically, ATAC-seq data was converted to a bedGraph reporting number of reads supporting a base position; then, these densities were converted to parts-per-10million by dividing each position’s density by a normalization factor based on the total library size. This library size-normalized data was used for downstream plotting

### Transposable element old versus young classification

To classify transposable elements (TEs) as old or young, we conducted a classification of EIE sequences listed in **Supplementary Table T2**. Elements were categorized based on their known evolutionary activity in humans. Young elements were defined as those from recently active subfamilies, including L1HS, L1PA2, SVA, and AluY, which are known to have current or recent retrotransposition activity in the human genome. Classifications can be found in **Supplementary Table 11.**

### CRISPR interference

The pHR-SFFV-dCas9-BFP-KRAB (Addgene, Cat# 46911) plasmid was modified to dCas9-BFP-KRAB-2A-Blast as previously described.^11^ Lentiviral particles were produced by co-transfecting HEK293T cells with the plasmid along with packaging plasmids psPAX2 and pMD2.G using a standard transfection method. Viral supernatants were harvested at 48 and 72 hours post-transfection, filtered through a 0.45 μm filter, and concentrated by ultracentrifugation at 25,000 rpm for 2 hours at 4°C. Cells were transduced with lentivirus, incubated for 2 days, selected with 1ug/ml blasticidin for 10–14 days, and BFP expression was analyzed by flow cytometry.

We took sgRNA specificity into account from the design phase of the CRISPRi screen. Our guide selection criteria included off-target scoring from Hsu et al. (2013)^11^ and filtering. We designed the library in benchling https://benchling.com with multiple independent sgRNAs per EIE element.

This redundancy helps distinguish on-target biological effects from off-target noise. To increase our stringency and ensure that the effects of low-efficiency or low-specificity guides do not interfere with the interpretation of the screen, we used FlashFry^12^ to score our gRNAs with multiple tools (**Supplementary Table 12**) and specifically selected the CRISPRi specificity score developed by Jost *et al.* 2020^13^ for filtering. We only report effects for elements with at least two guides that achieved a specificity score greater than 0.2, which is a standard cutoff for this type of scoring parameter (similar to the Doench *et al.* 2016^14^ CDF score).The oligo pool encoding guides (**Supplementary table T10**) were synthesized by Twist Bio and inserted into addgene Plasmid #52963 lentiGuide-Puro digested with Esp3I enzyme (NEB). The oligo pool was sequence validated. To investigate the effects of CRISPR interference, we utilized a lentiviral delivery system to introduce sgRNAs into cells stably expressing the dCas9-KRAB repressor.Lentiviral particles were produced as described above. The viral titer was determined by transducing HEK293T cells with serial dilutions of virus and assessing transduction efficiency via flow cytometry for GFP expression.

For transduction, cells were seeded at a density of 1 × 10^6 cells per well in 6-well plates and transduced overnight with lentivirus at a low multiplicity of infection (MOI) of 0.3, ensuring single sgRNA integration per cell. The following day, the medium was replaced with fresh growth medium. Two days post-transduction, cells were selected with 0.5 μg/mL puromycin for 4 days to enrich successfully transduced cells. GFP expression was monitored by flow cytometry to assess transduction efficiency. Post-selection, cells were harvested at multiple time points: baseline (day 4 after transduction), day 3, week 1, and month 1 (30 days). Genomic DNA was extracted using the DNeasy Blood & Tissue Kit (Qiagen) following the manufacturer’s instructions.

Integrated sgRNA sequences were amplified from genomic DNA using a multi-step PCR process. First, sgRNA cassettes were amplified using Primer set 1: hU6_pcr_out_fw (tggactatcatatgcttaccgtaacttgaaagt) and efs_pcr_rev (ctaggcaccggatcaattgccga). PCR reactions contained 0.8 μL each of 25 μM primers, 1-2 μg genomic DNA, water, and 25 μL NEB 2x master mix in a total volume of 50 μL. PCR conditions included an initial 3 min at 98°C, followed by 15-17 cycles of 20 s at 98°C, 20 s at 58°C, and 30 s at 72°C, concluding with a final extension for 1 min at 72°C. PCR products (∼400 bp) were verified by gel electrophoresis and purified. The second PCR step added Illumina sequencing adapters using primers (P5 stagger -hu6 and p7adpt_spRNAl105nt_rev). Reactions contained 10-50 ng purified PCR1 product, 0.8 μL each primer, water, and 25 μL NEB 2x master mix in 50 μL total volume, PCR 30 s at 98°C followed by 6 cycles of 15 s at 98°C, 15 s at 60°C, and 30 s at 72°C, finishing with 1 min at 72°C. PCR products (200-300 bp) were gel-verified and purified using AMPure XP beads. A final indexing PCR step was performed using Truseq-based P5 and P7 indexing primers. Reactions contained 10-50 ng DNA from PCR2, 0.8 μL each primer, water, and 25 μL NEB 2x master mix in 50 μL total volume. Conditions included 30 s at 98°C followed by 6 cycles of 15 s at 98°C, 15 s at 63°C, and 30 s at 72°C, ending with a 1-min extension at 72°C. Products were purified with AMPure XP beads and sequenced on an Illumina NextSeq platform using single-end 50 bp reads. Sequencing data were processed to quantify sgRNA representation at each time point, allowing analysis of sgRNA abundance dynamics over the experiment duration.

### CRISPRi fitness screen analysis

To compute the effect of each guide on cell fitness, we first quantified guide counts from sequencing libraries. To normalize counts across libraries, we converted raw guide counts to counts-per-million (CPM) and retained guides that had CPM values of at least 20 across all days tested. We also filtered out guides with high off-target scores (**Supplementary Table 12,** 0.2 cutoff from optimized CRISPRi design parameters^13^) and did not evaluate EIEs with <2guide after filtering. After confirming that normalized guide abundances were robust across replicates, we proceeded with our analysis using the average of guide replicates at each time point. We next scored the relative fitness of each guide against the non-targeting controls (NTC) by computing the ratio of CPM values between a guide and the NTC at the particular time point. Finally, we transformed this distribution to z-scores and reported this as the relative fitness effect of each guide.

### CRISPR-CATCH

In our study, we employed the CRISPR-CATCH (Cas9-Assisted Targeting of Chromosome segments) technique to isolate and analyze extrachromosomal DNA (ecDNA) structures. Following the standard protocol^15^, we designed two single-guide RNAs (sgRNAs) targeting specific enhancer regions: sgRNA #1 (ATATAGGACAGTATCAAGTA) and sgRNA #2 (TATATTATTAGTCTGCTGAA). These sgRNAs directed the Cas9 nuclease to introduce double-strand breaks at the targeted sites, linearizing the circular ecDNA molecules. The linearized DNA was then subjected to pulsed-field gel electrophoresis (PFGE) using S. cerevisiae and H. wingei DNA ladders as molecular weight markers to facilitate size-based separation. Distinct DNA bands corresponding to the targeted ecDNA were excised from the gel for downstream analyses, including sequencing.

### Probe Design

Probes were designed against human genome assembly hg19, tiling the regions in **Supplemental Table T7** using the probe designing software described previously.^16,17^ We restricted choice of the 40mer targeting region of the probes to a GC range of 20-80%, a melting temperature of 65-90 degrees centigrade, and excluded sequences with non-unique homology (cut off of 17mer homology to any other sequence in the genome) or with homology to common repetitive elements in the human genome listed in repbase (cut off of 14mer). Targeting probes were then appended with a 20mer barcode per target region. Probe design software is available at https://github.com/BoettigerLab/ORCA-public. Finalized probe libraries were ordered as an oligo-pool from Genscript.

### ORCA imaging

ORCA hybridization was performed as previously described.^17,18^ Briefly, 40mm Bioptechs coverslips were prepared with EMD Millipore™ Poly-D-Lysine Solution (1 mg/mL, 20mL, dilute 1:10)(Sigma, cat. No. A003E) for 40 minutes. Coverslips were then rinsed 3x in 1x PBS. Cells were passaged onto the coverslips and allowed to adhere overnight. The next day, the coverslip with cells were rinsed 3 times in 1x PBS and then fixed for 10 minutes in 4% PFA. For DNA imaging: Cells were then permeabilized in 0.5% Triton-x 1x PBS for 10 minutes followed by 5 minutes of denaturing in 0.1M HCL. A 35-minute incubation in hybridization buffer prepared samples for the primary probe. Primary probes were added (1ug) directly to the sample in hybridization solution and then the sample was heated to 90 degrees celsius for 3 minutes. An overnight 42-degree incubation (or at least 8 hour incubation) was performed followed by post-fixation in 8% PFA + 2% glutaraldehyde in 1× PBS before being stored in 2x SSC or used immediately for imaging. For RNA imaging, the HCL, heat, and post-fixation steps were omitted.

DNA samples were imaged on one of two different homebuilt setups designed for ORCA, “scope-1”, “scope-3”, depending on instrument availability. Microscope design parameters were deposited in the Micro-Meta App.^19^ The design and assembly of the “scope-1” system is described in detail in our prior protocol paper.^20^ Both systems use a similar auto-focus system, fluidics system, and sCMOS camera (Hamamatsu FLASH 4.0), though scope-3 had a larger field of view (2048×2048 108 nm pixels) compared to scope-1 (1024×1024 154 nm pixels).

RNA samples were imaged on a different homebuilt setup designed for ORCA designated as the “Yale lumencor system”. This system uses a similar auto-focus system and fluidics system, with a sCMOS camera (Hamamatsu ORCA BT fusion) with a field of view (2304×2304 108nm pixels) and Olympus PlanApo 60x objective .

Automated fluidics handling is described in detail in our prior protocol paper.^17^ Briefly, fluid exchange between each imaging step was performed by a homebuilt robotic setup. The system used a 3-axis CNC router engraver, buffer reservoirs and hybridization wells (96-well plate) on the 3-axis stage, ETFE tubing, imaging chamber (FCS2, Bioptechs), a needle, and peristaltic pump (Gilson F155006). The needle was moved between buffers or hybridization wells and was flown across the samples through tubing using the peristaltic pump. Open-source software for the control of the fluidics system is described in the “Software Availability” section below.

Sequential imaging of ORCA probes was conducted alternating between hybridization of fluorescent adapter probes, readout probes complementary to the barcodes on the primary probe sequences, imaging, and stripping of probes, as described previously.^17,18^ Briefly, a z-stack was acquired over 10um at 250nm step size where each step alternated lasers between data channel and fiducial. Readout probes were labeled with Alexa-750 fluorophores. Fiducial probe was labeled in cy3 and added only in the initial round. RNA imaging was performed with the EIE 14 probe labeled with the Alexa-750 and the *MYC* probe labeled with the Cy5 fluorophores.

Sequence for the fiducial: /5Cy3/AGCTGATCGTGGCGTTGATGCCGGGTCGAT

Sequence of Cy5: /5Cy5/TGGGACGGTTCCAATCGGATC

Sequence of the 750:/5Alex750N/ACCTCCGTTAGACCCGTCAG

### Image processing

Image processing was performed with custom MATLAB functions available: https://github.com/BoettigerLab/ORCA-public. Briefly, cells were max projected and pixel-scale alignment was computed across all fields of view off of the fiducial signal. This alignment was then applied in 3D across all 250 nm z steps. Cellpose^21^ was then used to segment individual cells. A cell-by-cell fine scale (subpixel) alignment was then computed and aligned individual cells were then ready for 3D-spot calling. The individual ecDNA spots and their 3D positions computed to sub-pixel accuracy using the corresponding raw 3D image stacks and the 3D DaoSTORM function in storm-analysis toolbox [DOI: 10.5281/zenodo.3528330] an open source software for single-molecule localization, adapted for dense and overlapping emitters following the DaoSTORM algorithm.^22^ DaoSTORM was run in the 2d-fixed mode, as the 3D fitting modes are for estimating axial position from astigmatism in the xy plane, rather than computing it directly from a z-stack. The fixed-width PSF of the microscope is pre-computed using 100 nm (sub-diffraction) fluorescent beads. A minimum detection threshold of 30 sigma was used for the fit. The z-position of the localizations was computed using Gaussian fit to the vertically stacked localizations, with an axial Gaussian width also pre-computed from z-stack images with 100 nm fluorescent beads. Additional information can be found in the read-the-docs for storm-analysis: https://storm-analysis.readthedocs.io/en/latest/.

### Minimum pairwise distance quantification

All pairwise distances between genomic regions were calculated on a per-cell basis. The shortest distances were saved for each *MYC* centroid and EIE 14 and *PVT1* such that each *MYC* centroid has one corresponding shortest distance per EIE 14 and *PVT1*. For each cell, a sphere radius *r=*4um (the average radius of cells calculated with Cellpose mask) with randomly simulated points corresponding to the number of *MYC*, EIE 14, and *PVT1* centroids. The same minimum pairwise distance quantification was calculated on the randomly simulated points.

### Ripley’s K quantification

To calculate the density corrected distance ratios a distance cutoff of 2um and an interval density of 0.01:0.01:2 was used. The spatial relationship between *MYC* and EIE 14 and *MYC* and *PVT1* were quantified as follows: On a per-cell basis the distance density function was calculated, truncated at the specified cutoff. A uniform distribution was then computed over the same interval and a ratio of these values was taken. This ratio was then corrected by the volume of the interval shell.

### Reporter plasmid construction and transfection

All plasmids are made with Gibson assembly (NEB HIFI DNA assembly kit) according to manufacturer’s protocol. We used a plasmid from this publication^9^ containing the *MYC* promoter (chr8:128,745,990–128,748,526, hg19) driving NanoLuc luciferase (PVT1p-nLuc) and a constitutive thymidine kinase (TK) promoter driving Firefly luciferase, this plasmid was used as negative control. pGL4-tk-luc2 (Promega) plasmids with an enhancer (chr8:128347148– 128348310) was used as positive control.^9^ In the test plasmid, the *cis*-enhancer was replaced by 1.7 kb sequence of EIE 14 or by Part #1: L1PA2 or by Part #2: L1M4a1 (**Supplementary Table T13**). To assess luciferase reporter expression, COLO320DM cells were seeded into a 24-well plate with 100,000 cells per well. Reporter plasmids were transfected into cells the next day with lipofectamine 3000 following the manufacturer’s protocol, using 0.25 μg DNA per well. Luciferase levels were quantified using Nano-Glo Dual reporter luciferase assay (Promega).

### Statistics and reproducibility

All statistical tests used, replicate information, and sample size information are reported in the figure legends. No statistical method was used to predetermine sample size. No samples or data points were excluded. The experiments were not randomized. The investigators were not blinded to the conditions of the experiments during data analysis.

## Data availability

All sequencing data generated in this study is available through the Gene Expression Omnibus (GEO) accession number GSE277492.

https://www.ncbi.nlm.nih.gov/geo/query/acc.cgi?acc=GSE277492 and BioProject NCBI ID: 1162466. https://www.ncbi.nlm.nih.gov/bioproject/1162466

Raw RNA imaging data related to figure 5 is hosted here: https://doi.org/10.5281/zenodo.16921322

All raw imaging data related to the DNA is available upon request as it is large. The processed data tables from image analysis recording x,y,z positions of RNA and DNA can be found: https://github.com/sedona-Eve/Kraft_Murphy_Jones_ecDNA/

## Code Availability

The image analysis code is publicly available at: https://github.com/BoettigerLab/ORCA-public/ and https://storm-analysis.readthedocs.io/en/latest/analysis.html. Code for reconstructing amplicons from long read data with the CoRAL algorithm is also publicly available: https://github.com/AmpliconSuite/CoRAL

---

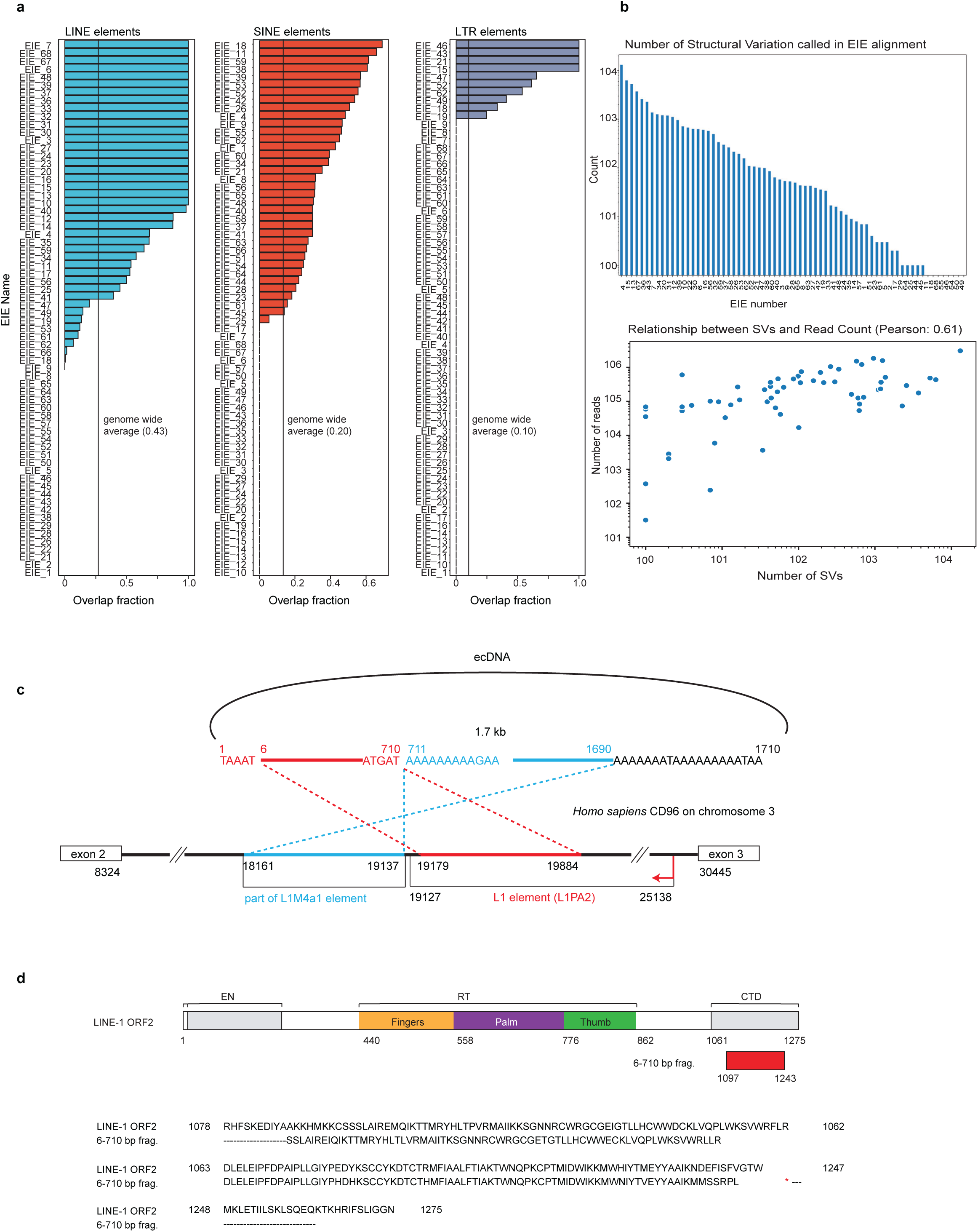

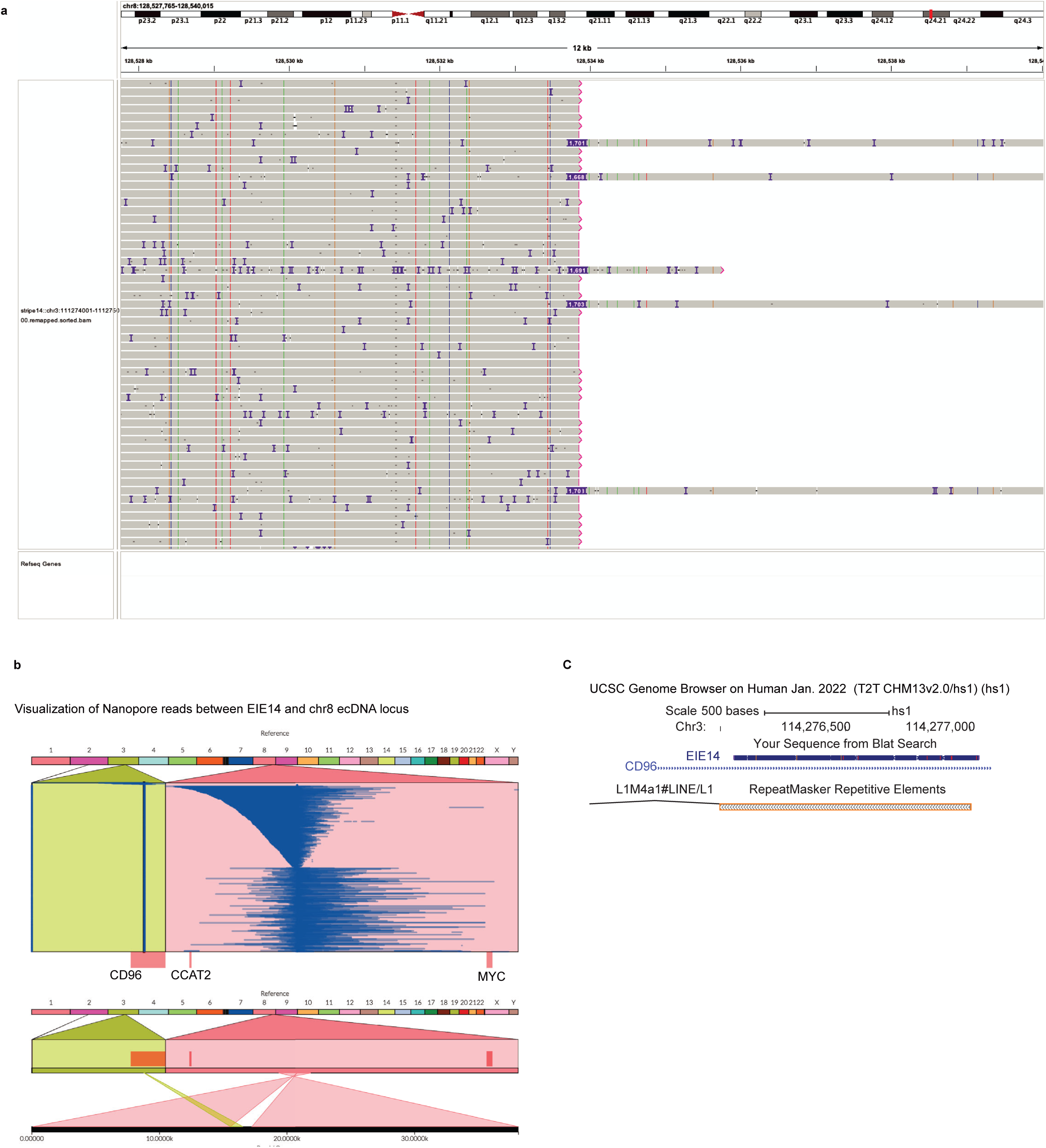

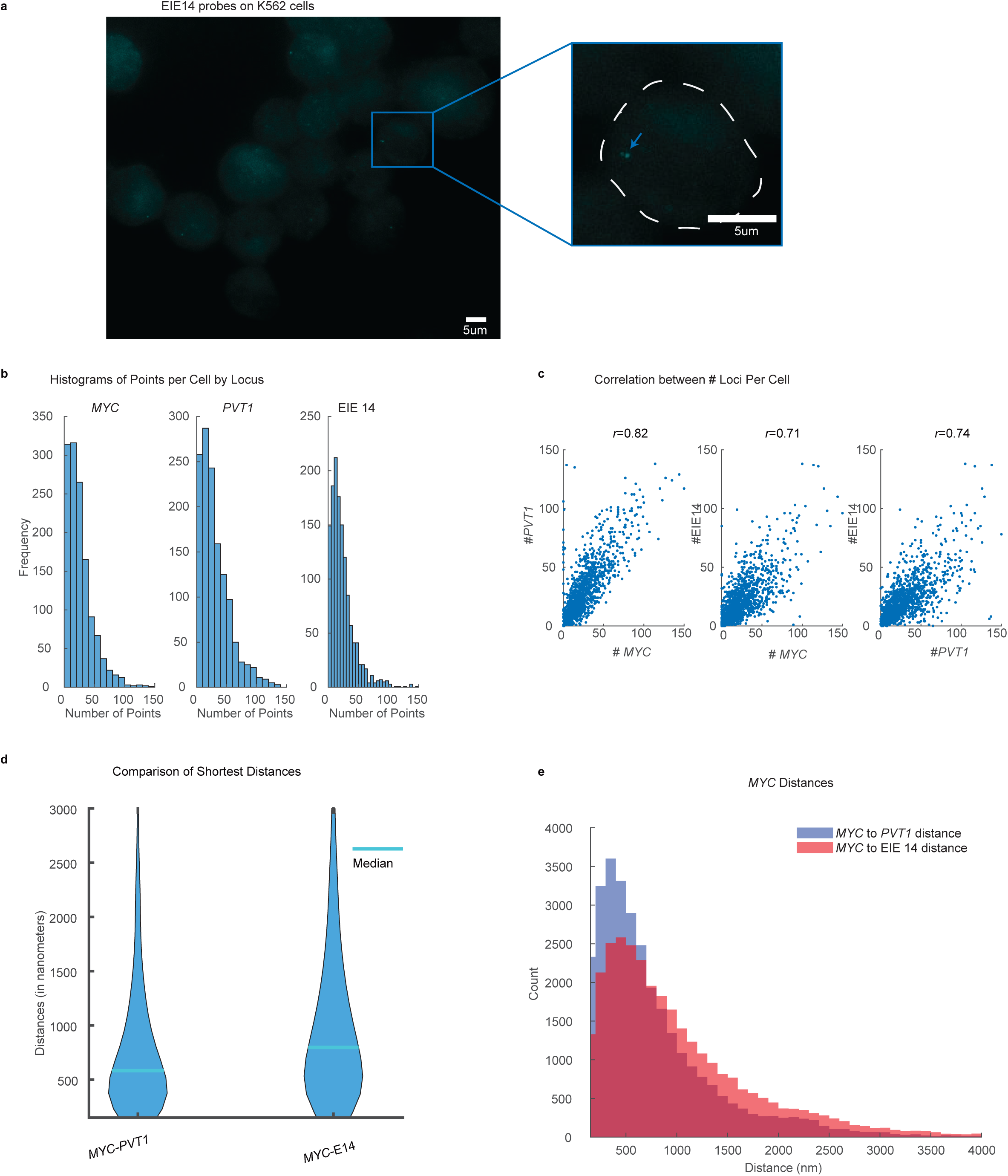

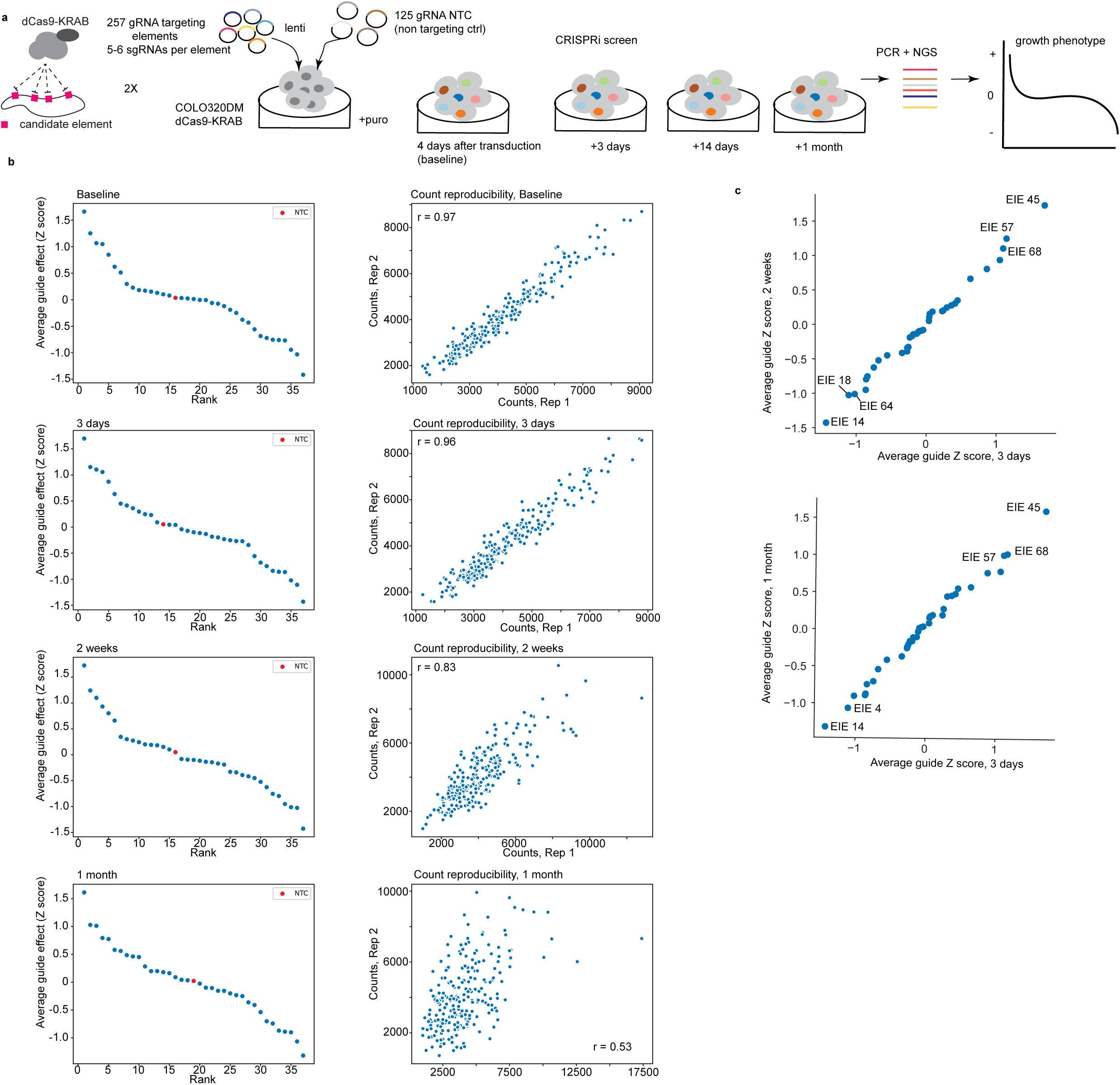

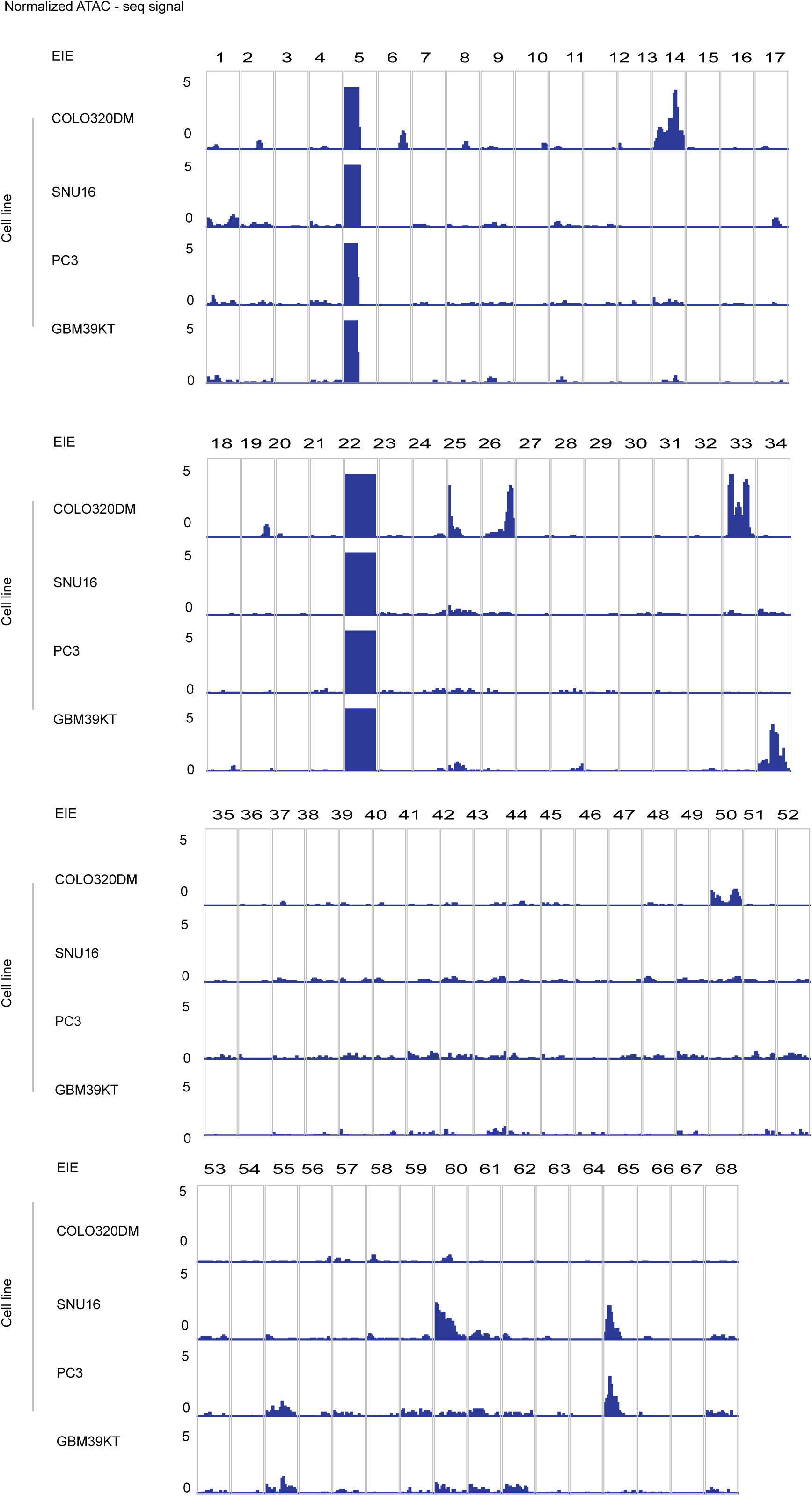

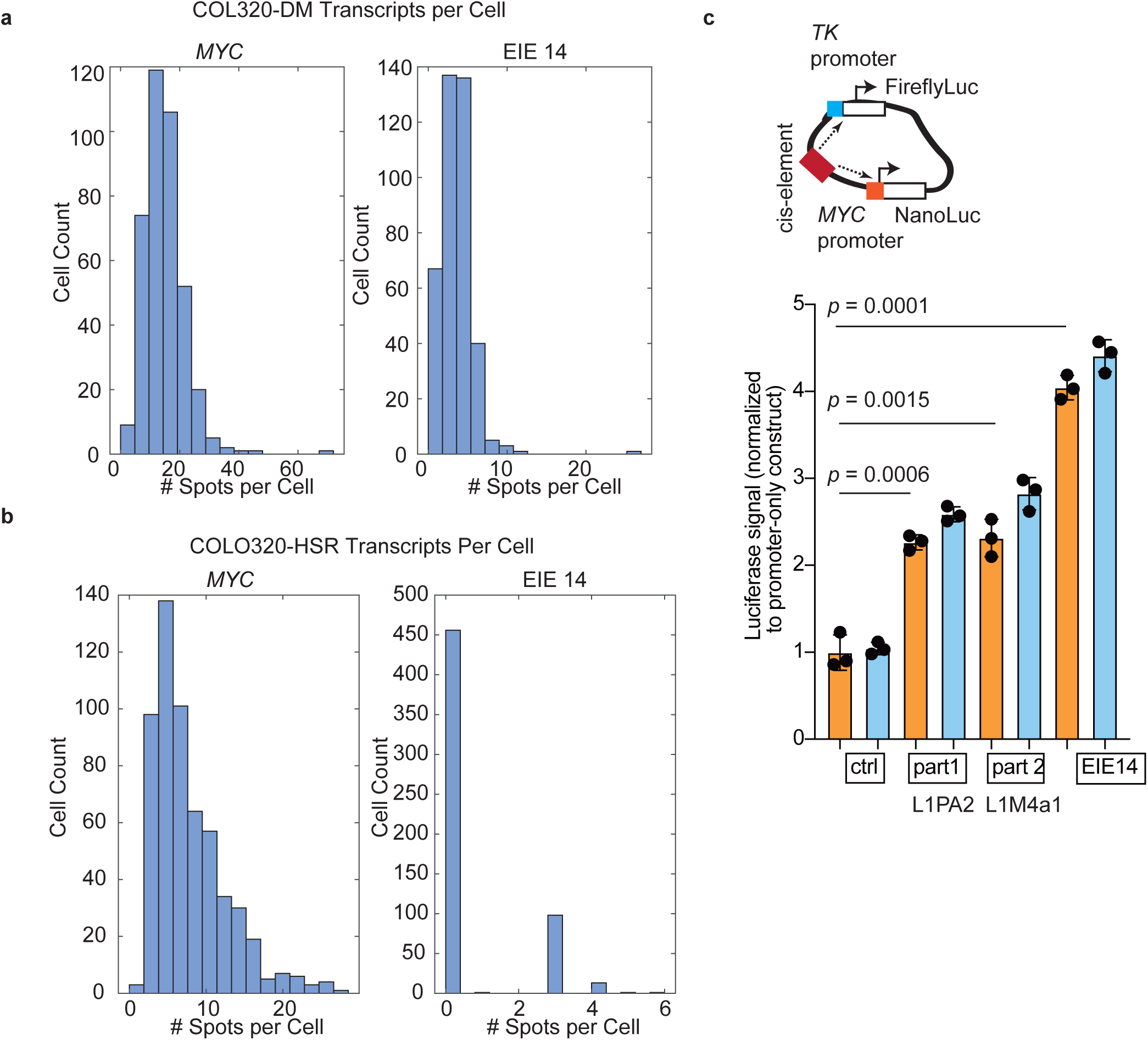

